# Ultrastructural remodelling of tau fibrils during ghost tangle formation in Alzheimer’s disease brain

**DOI:** 10.64898/2026.07.21.739778

**Authors:** Daniel A. Stähli, Louis Travers, Notash Shafiei, Lukas van den Heuvel, Eglantine Vialaneix, Philippine L. Schneider, Annemieke J.M. Rozemuller, Wilma D. J. Van de Berg, Henning Stahlberg, Amanda J. Lewis

**Affiliations:** Laboratory of Biological Electron Microscopy, Institute of Physics, School of Basic sciences, EPFL, Lausanne, Switzerland; Department of Fundamental Microbiology, Faculty of Biology and Medicine, University of Lausanne, Switzerland; Department Anatomy and Neurosciences, section Clinical Neuroanatomy and Biobanking, Amsterdam University Medical Centre, Vrije University, Amsterdam, The Netherlands; Amsterdam Neuroscience, program Neurodegeneration, Amsterdam University Medical Centre, Vrije University, Amsterdam, The Netherlands

**Keywords:** Alzheimer’s disease, correlative light and electron microscopy, disease pathology, post-mortem human brain, neurofibrillary tau tangles, astrocytes

## Abstract

Tau aggregation into intracellular neurofibrillary tangles (NFTs) is one of the major hallmarks of Alzheimer’s disease (AD). Based on neuropathological studies, NFTs have been classified into pre-tangles, mature tangles, and ghost tangles, however the ultrastructural transitions between these stages remain poorly understood. Here, we used correlative light and electron microscopy (CLEM) to structurally characterize tau tangle maturity states in post-mortem human AD brain tissue. Pre-tangles showed no consistent fibrillar ultrastructure. Mature tangles contained densely packed, highly aligned paired helical filaments (PHF) and straight filaments (SF), often organized in spatially distinct bundles within the neuronal soma. Ghost tangles lacked cellular organelles and were composed predominantly of thin fibrils compartmentalized by membranous structures, with fibril morphology differing between compartmentalized and non-compartmentalized regions. Electron tomography and fibril segmentation demonstrated that these fibrils were significantly thinner than PHFs and SFs while immunogold labeling using the 2E9 tau marker confirmed the presence of tau within both mature and ghost tangle fibrils. GFAP-positive astrocytic processes infiltrated fibril-rich compartments within ghost tangles, linking astrocytic engagement with the emergence of this distinct ultrastructural organization. Together, our findings show that ghost-tangles contain a structurally distinct population of tau fibrils, suggesting that tau aggregates undergo astrocytic-mediated structural remodeling at late stages of pathology.

## Introduction

Alzheimer’s disease (AD) is a progressive neurodegenerative disease. It remains the predominant driver of dementia worldwide^1^. Neuropathologically, AD is classically defined by two hallmark protein aggregates in the brain: extracellular amyloid beta (Aβ) plaques and intracellular neurofibrillary tangles (NFTs) formed by hyperphosphorylated tau (p-tau)^2^. Despite decades of research, no cure exists, and most current treatments are symptomatic rather than disease-modifying^1,3,4^. Imaging studies have shown that p-tau pathology correlates more closely than Aβ plaques with neuronal loss and cognitive decline^5,6^. Consequently, tau has emerged as a promising therapeutic target^3^, highlighting the importance of understanding NFT formation and maturation.

Recent neuropathological studies by Moloney et al. have classified NFTs into three maturity levels: pre-tangles, mature tangles, and ghost tangles based on immunohistochemical staining^7,8^. Besides NFTs, pathological p-tau aggregation can also be detected in swollen dystrophic neurites, thinner neuropil threads or astrocytes^9^. Electron microscopy (EM) described the fibrillar ultrastructure of mature tangles and neuritic pathology composed of aligned twisting paired helical filaments (PHFs) and straight filaments (SFs)^10–13^, whereas pre-tangles appear to be mostly devoid of fibrils^9^. Dystrophic neurites are also associated with the presence of membranous material called dense bodies or multilamellar bodies^11,14^. Ghost tangles are described to be extracellular, lack a nucleus^7^, and are frequently associated with astrocytic processes and membranous compartments^13,15–18^. Thin, non-twisting filaments have been reported in ghost tangles^13^, but their molecular identity and relationship to PHFs and SFs remain unclear.

The atomic structure of PHFs and SFs has been described by cryo-EM from tau fibrils extracted from AD brains^19,20^, and in situ cryo-electron tomography has confirmed the PHF fold within human tissue^21^. The presence of multiple different fibril types in ghost tangles, the majority being much thinner than PHFs, raises the question of their identity, especially given that ghost tangles are often infiltrated by astrocytes^13,15–18^. Until now, a systematic ultrastructural comparison of the different stages of tau pathology in the human brain has not yet been performed, and the structural nature of fibrils specifically within ghost tangles remains incompletely defined.

Defining the mechanisms underlying tau tangle maturation and identifying when and how these processes can be therapeutically targeted, is critical to advance disease-modifying intervention strategies. Here, we used a correlative light and electron microscopy (CLEM) pipeline^22^ to identify and characterize tau pathology across defined maturity stages in the hippocampus and entorhinal cortex from post-mortem AD donors. By combining immunofluorescence-based staging with ultrastructural analysis, electron tomography, and immunogold labelling, we aimed to define how tau fibril architecture changes across NFT maturation and to determine the ultrastructural nature and molecular identity of fibrils within ghost tangles. We showed that tau fibrils differ in morphology across NFT maturity stages, with ghost tangles containing a distinct population of thinner tau-positive filaments.

## Results

### Identification of tau tangle maturation classes

We included clinically defined and pathologically confirmed AD cases with severe NFT pathology in the hippocampus and entorhinal cortex from four brain donors (**Suppl. Table 1**). Free-floating vibratome sections (20 – 60 µm thick) were immunolabelled to localize pathology using immunofluorescence microscopy. Antigen retrieval was omitted from the immunolabelling to preserve the tissue ultrastructure for the downstream EM. To address potential issues with antigenicity or antibody penetration, we tested a panel of tau antibodies targeting different phosphorylation sites, peptide sequences, C-terminal truncation and a conformational tau marker (**Suppl. Table 2**, **Suppl. Fig. 1**). We also tested the β- sheet marker Amytracker and two granulovacuolar degeneration markers, which have been described to be present in neurons with pre-tangles^23^. Eight out of nine tested p-tau markers were found to label NFT pathology efficiently, while the granulovacuolar degeneration markers and some tau markers for later tangle maturity levels did not produce strong, consistent labeling in our hands (**Suppl. Table 2**), possibly due to the detergent-free immunolabeling procedure that is necessary to preserve tissue ultrastructure. We further tested several of these markers for immunohistochemistry (IHC) on 200 nm resin-embedded sections and immunogold labeling directly on 80 nm sections on EM grids and identified antigen retrieval conditions under which the tested markers showed efficient labeling (**Suppl. Table 2**).

The final fluorescent labelling of the free-floating sections for the identification and localization of the different tangle maturation stages included a marker for p-tau (pT217, AT8, or AT180), a marker for DNA (DAPI), and Amytracker 480 (**Suppl. Table 2**). Tau tangle maturity levels were assessed based on the presence or absence of these markers and the characterization scheme previously defined by Moloney et al^8^. Pre-tangles were characterized by weak staining for p-tau, together with the presence of a nucleus and the absence of strong β-sheet (Amytracker) staining. Mature tangles were positive for all markers (tau, p-tau and Amytracker), and ghost tangles showed strong β-sheet staining (Amytracker) but no p-tau positivity and had no associated nucleus (**Suppl. Fig. 2**). Using our previously described CLEM workflow^22^, we targeted and analyzed the ultrastructure of 8 pre-tangles, 43 mature tangles, 41 ghost tangles (**Suppl. Table 3**), as well as neuritic pathologies, of which representative images are shown in this manuscript. Some degree of ultrastructural deterioration in the tissue was observed (for example empty extracellular space or cytoplasm, and discontinuous plasma membranes^24^), which may reflect a combination of advanced disease stage, prolonged post-mortem delay and tissue processing artefacts.

### Pre-tangle-bearing neurons are ultrastructurally identical to neighboring tau immunonegative cells

Pre-tangles showed weak diffuse fluorescent signal for multiple p-tau antibodies (**Suppl. Table 2**), presence of a nucleus and lack of Amytracker staining in the free-floating sections. Additionally, weak dotted cytoplasmic immunostaining, often with marked perinuclear staining, was observed in the ultra-thin sections (**Suppl. Fig. 2**). We did not detect any consistent ultrastructural differences in the cellular morphology of pre-tangle-bearing neurons and neighboring tau-immunonegative cells (as confirmed by the absence of tau/amytracker staining) (**Suppl. Fig. 3**). Lipofuscin granules, which are pigmented or granular lipid droplets and a normal feature of neurons in the hippocampus and entorhinal cortex in the ageing brain^25–29^ were observed. Cellular organelles such as endoplasmic reticulum (ER), the Golgi apparatus and mitochondria appeared intact when visible. Electron-dense vesicles, consistent with the ultrastructure described for lysosomes, were abundant and displayed a typical morphology (approximately 0.5µm in diameter), and autophagosomes were occasionally observed. Some pre-tangles also contained vacuoles that may represent granulovacuolar bodies reported to be a hallmark of early AD pathology^30,31^. However, well defined granulovacuolar bodies containing dense cores were only observed once (**Suppl. Fig. 3B**).

In one pre-tangle, perinuclear fibrils were observed (**Suppl. Fig. 3C**). It is possible that this may represent an intermediate state between pre-tangles and mature tangles; however, similar fibril-like structures were also found in tau-immunonegative cells (**Suppl. Fig. 3 D1, D2**). Therefore, it is likely that these perinuclear fibrils represent cytoskeleton filaments.

### Mature tangles contain highly organized tau fibril bundles and intact organelles

Mature tangles were positive for tau, p-tau and Amytracker, and were consistently associated with a nucleus, supporting their intracellular localization. The ultrastructure of mature tangles often showed large quantities of tau fibrils, which were identified by their morphology (typical thickness, clearly visible twist), by the presence of tau-immunopositive staining on adjacent IHC sections, and/or by their correlation with a fluorescence signal of the free-floating section (**Fig. 1**). The plasma membrane of the cell was visible and cell organelles such as ER, Golgi apparatus and mitochondria appeared intact compared to neighboring tau-immunonegative cells (**Fig. 1B,C**, **Suppl. Fig. 4C,D** and **Suppl. Fig. 5A1, A2**). Lysosomes and autophagosomes were also observed; however, lysosomes were noticeably less abundant than in pre-tangles. When present, they exhibited a more granular ultrastructure compared to those observed in pre-tangles (**Suppl. Fig. 4D**).

**Figure 1:**
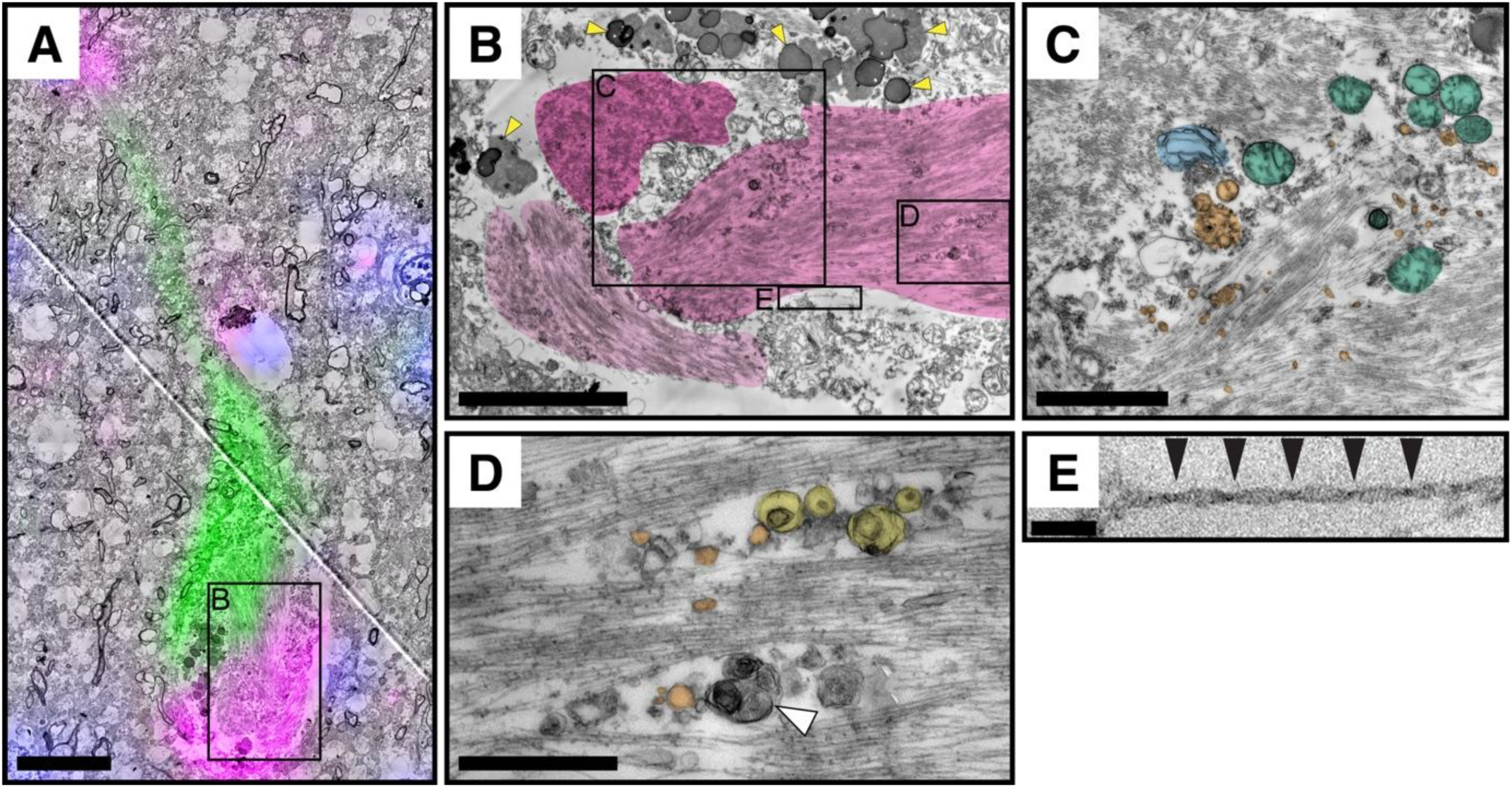
Ultrastructural characterization of a mature tangle. **A** Correlative fluorescence microscopy showing phosphorylated tau (pT217, magenta), β-sheet signal (Amytracker, green) and nuclei (DAPI, blue) aligned with the corresponding EM micrograph or a representative mature tangle. **B,** The neuronal soma is filled with densely packed tau fibrils organized into spatially distinct bundles with different orientations (pseudo-colored). Lipofuscin is visible above the fibril-rich regions (yellow arrowheads). **C,** Mitochondria (green), vesicles (orange) and Golgi/ER-like membranes (blue) can be present between fibril bundles. **D,** multi-vesicular (yellow) or multilamellar (white arrowhead) structures occasionally occur within fibril bundles. **E,** High-magnification view of one of the PHFs showing the characteristic periodic twist (arrowheads indicate constrictions). Scale bars: a = 10 µm; b = 5 µm; c = 2 µm; d = 1 µm; e = 100 nm.

Lipofuscin was frequently observed in cells containing mature tangles (**Fig. 1b**), as well as multilamellar bodies (**Suppl. Fig. 5A**). Notably, while such lipidic/membranous material was observed within or adjacent to fibril-rich regions, it appeared spatially segregated from the tau fibril bundles.

Most mature tangles contained large amounts of fibrils that appeared to be organized in bundles (**Fig. 1B-D**, **Suppl. Fig. 5A**), whereas some contained relatively few tau fibrils, in spatially distinct clusters (**Suppl. Fig. 4**), Other cellular organelles and vesicular material was occasionally present but appeared segregated from fibril bundles (**Fig. 5C,D**).

Dystrophic neurites, which were abundant in the tissue, are typically long and thin and therefore difficult to trace across adjacent ultrathin sections. For this reason, they were predominantly identified in the EM by their morphology alone and not through correlation to IHC and fluorescence images. Dystrophic neurites contained twisted fibrils and appeared swollen in comparison to neighboring neuropil threads and other cellular processes lacking these characteristics (**Suppl. Fig. 5**). Additionally, dystrophic neurites showed ultrastructural heterogeneity ranging from predominantly membranous, containing large quantities of multilamellar bodies and other vesicular material, to predominantly fibrillar morphologies (**Suppl. Fig. 5**). Both ultrastructural variants were tau immuno-positive where correlation was possible (data not shown).

### Ghost tangles are compartmentalized and contain highly organized fibrils and vesicular material

Ghost tangles lacked p-tau immunoreactivity but were positive for the tau antibody 2E9, whose epitope maps to the core of tau fibrils (**Suppl. Fig. 1**), and Amytracker (**Suppl. Fig. 2**). By EM, ghost tangles lacked a discernible plasma membrane, nucleus or intact cell organelles, in agreement with the literature^13,17,18^, and confirming their neuronal disintegration and degenerative features. Of note, they were composed of compartmentalized fibrillar regions delineated by membrane structures (**Fig. 2B,C**). The fibrils in ghost tangles appeared to be thinner, more densely packed, and lacked a visible twist (**Fig. 2C**) compared to PHFs and SFs observed in mature tangles. Between the compartmentalized fibrils, some vesicular material and mitochondria were sometimes present, but no other cellular organelles (ER/Golgi) or lipofuscin were observed (**Fig. 2B**). The compartmental organization of thin fibrils was observed across several CLEM cycles of the same ghost tangle, in total spanning several μm of tissue, representing the majority of fibrillar material comprising ghost tangles (**Suppl. Fig. 6**).

**Figure 2:**
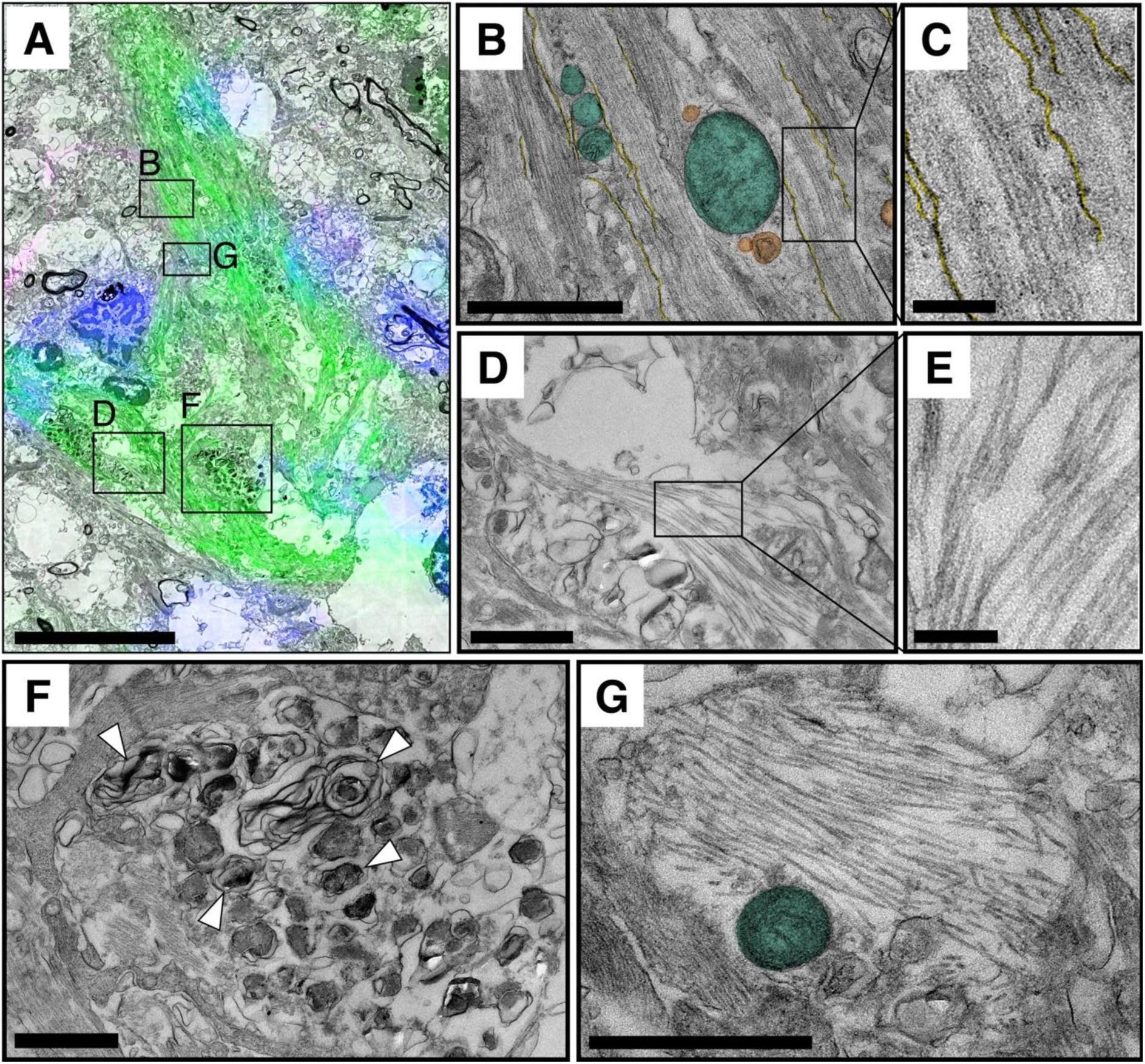
Ultrastructural characterization of a ghost tangle. **A,** Correlative fluorescence microscopy showing β-sheet signal (Amytracker, green), an absence of phosphorylated tau (pT217, magenta) or an associated nucleus (DAPI), correlated with the corresponding EM micrograph of a representative ghost tangle. Surrounding nuclei belong to neighboring cells. **B,C,** Compartments containing thin, densely packed fibrils lacking a visible twist are delineated by membrane profiles (yellow). Vesicles (orange) and a mitochondrion (green) are present within some compartments. **D,E,** Occasional PHFs/SFs are observed adjacent to, but outside, the thin-fibril-rich compartments. **F,** Within the ghost tangle there also appears to be a dystrophic neurite containing multilamellar bodies and **G,** a neuropil thread containing straight filaments. Scale bars: A = 10 µm; B = 1 µm; C = 200 nm; D = 2 µm; E = 200 nm; F = 1 µm; G = 1 µm.

PHFs and SF were occasionally observed close to ghost tangles, typically outside the thin-fibril-rich compartments (**Fig. 2D,E**). The PHFs/SFs were sometimes in regions that were not delineated by a membrane, appearing to be in the extracellular space (**Fig. 2D,E**).

Ghost tangles could also contain other distinctive compartments, including dystrophic neurites, known as tangle associated neuritic clusters (TANCs)^7,32^, like those observed in neuritic Aβ plaques^12,14,18^, and contained multiple multilamellar bodies (**Fig. 2F**), as well as neuropil threads containing SFs (**Fig. 2G**).

### Ghost tangle fibrils contain an ultrastructurally distinct class of tau fibrils

Based on our CLEM data, fibrils within ghost tangles appeared structurally distinct from tau fibrils in mature tangles. To quantify this distinction, we performed electron tomography and fibril segmentation on fibrils from within one mature tangle and one ghost tangle.

Additionally, as astrocytes have been shown to infiltrate ghost tangles^16–18,33–35^, we analyzed fibrils from within a putative astrocytic process that resembled the known GFAP filament ultrastructure, based on their thin diameter, lack of twisting, dense packing, and high curvature. We later validated the ultrastructural identity of similar GFAP-like fibrils by CLEM on adjacent tissue sections (**Suppl. Fig. 7**).

This analysis showed PHFs in mature tangles to measure ∼23 nm diameter at their widest area, ghost tangle fibrils ∼16 nm, and the putative GFAP fibrils ∼14 nm (**Fig. 3**). Measured widths differed from some previously reported values (∼25 nm for PHF and 16-20 nm for SFs^12,17,18,36^, 8-9 nm for GFAP^37^), which may reflect differences in preparation and analysis methods. Nevertheless, all three fibril classes showed significantly different width distributions (p<0.005), with the fibrils in ghost tangles being thinner than PFs and SFs in mature tangle fibrils and thicker than GFAP fibrils, confirming our ultrastructural observations.

**Figure 3:**
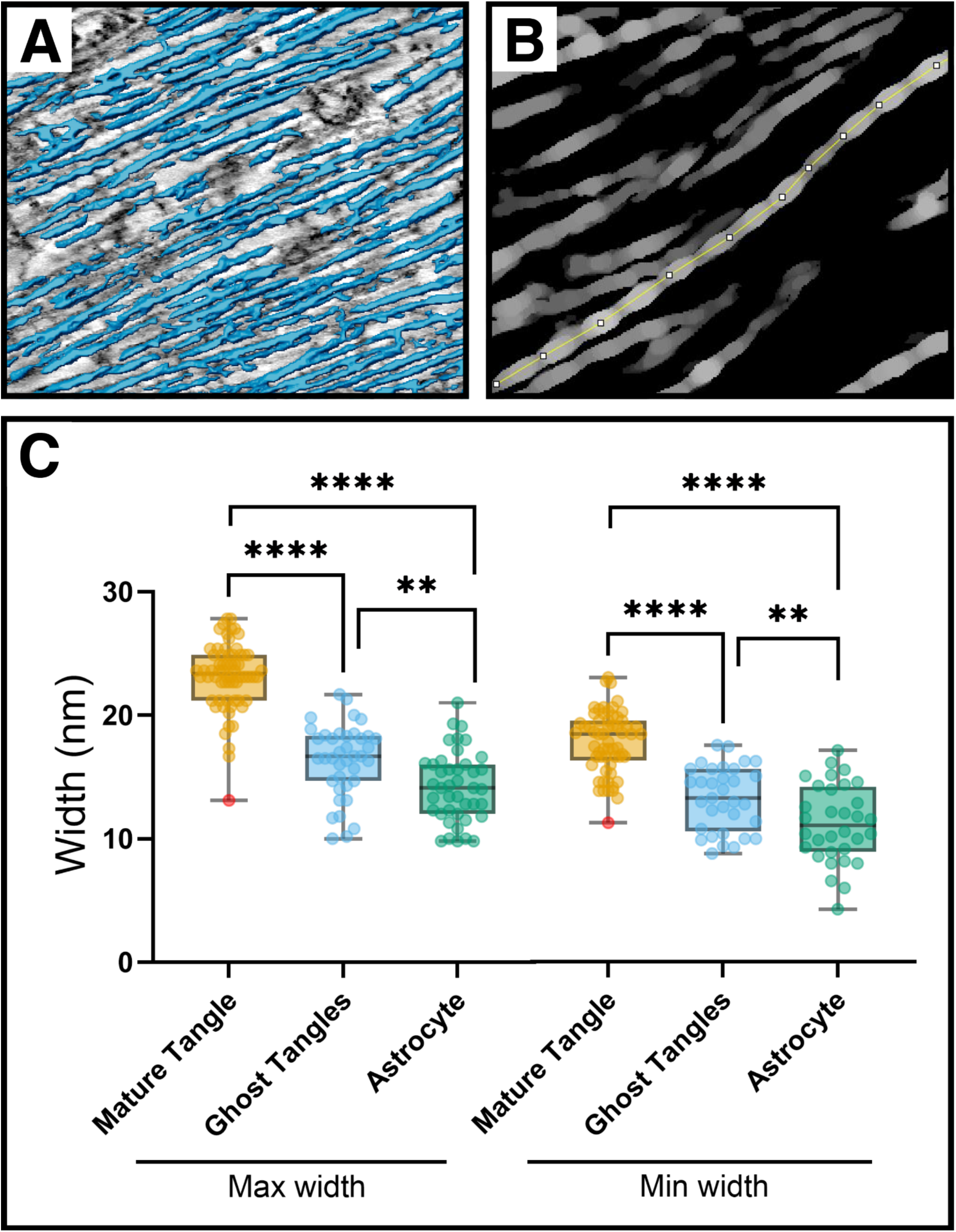
Fibrils in ghost tangles are ultrastructurally distinct from PHFs and SFs found in mature tangles. **A,** An electron tomogram of a mature tangle with segmented PHFs overlayed. **B,** thickness map of an individual PHF showing measurement of local maxima and minima across the fibril width. **C,** Quantification of fibril widths from mature tangles (PHFs/SFs), ghost tangles (thin tau fibrils) and astrocytic processes (GFAP filaments). Each point represents one measurement. The widths of all three fibril types differed significantly (**= p<0.001, ****= p<0.0001), as calculated by Welch’s-test. Outliers were identified using Z-scores < −3 or > 3 and are indicated in red.

To investigate the molecular identity of the thinner ghost tangle fibrils, we performed immunogold labeling with GFAP and the tau antibody 2E9, which detects both mature and ghost tangles. Labelling with 2E9 showed enrichment of gold particles over fibrillar regions in both the mature tangle and ghost tangles (**Fig. 4A-C Suppl. Fig. 8**) whereas local vesicles and mitochondria were not labeled. In the ghost tangles, the 2E9 labeling was associated with the thin fibrillar compartments and SFs in their vicinity (**Fig 4 A-C**), indicating that the thin fibrils within ghost tangles are tau-immunopositive. GFAP immunogold labeling was detected in astrocytic processes adjacent to ghost tangles (F**ig 4D,F**) and was absent from mature tangles, with GFAP-positive processes restricted to the surrounding neuropil (**Suppl. Fig. 8**), confirming labelling specificity. GFAP labeling was also detected in a ghost tangle (**Fig 4E,D**), consistent with previously reported astrocytic infiltration of ghost tangles^16–18,33–35^.

**Figure 4:**
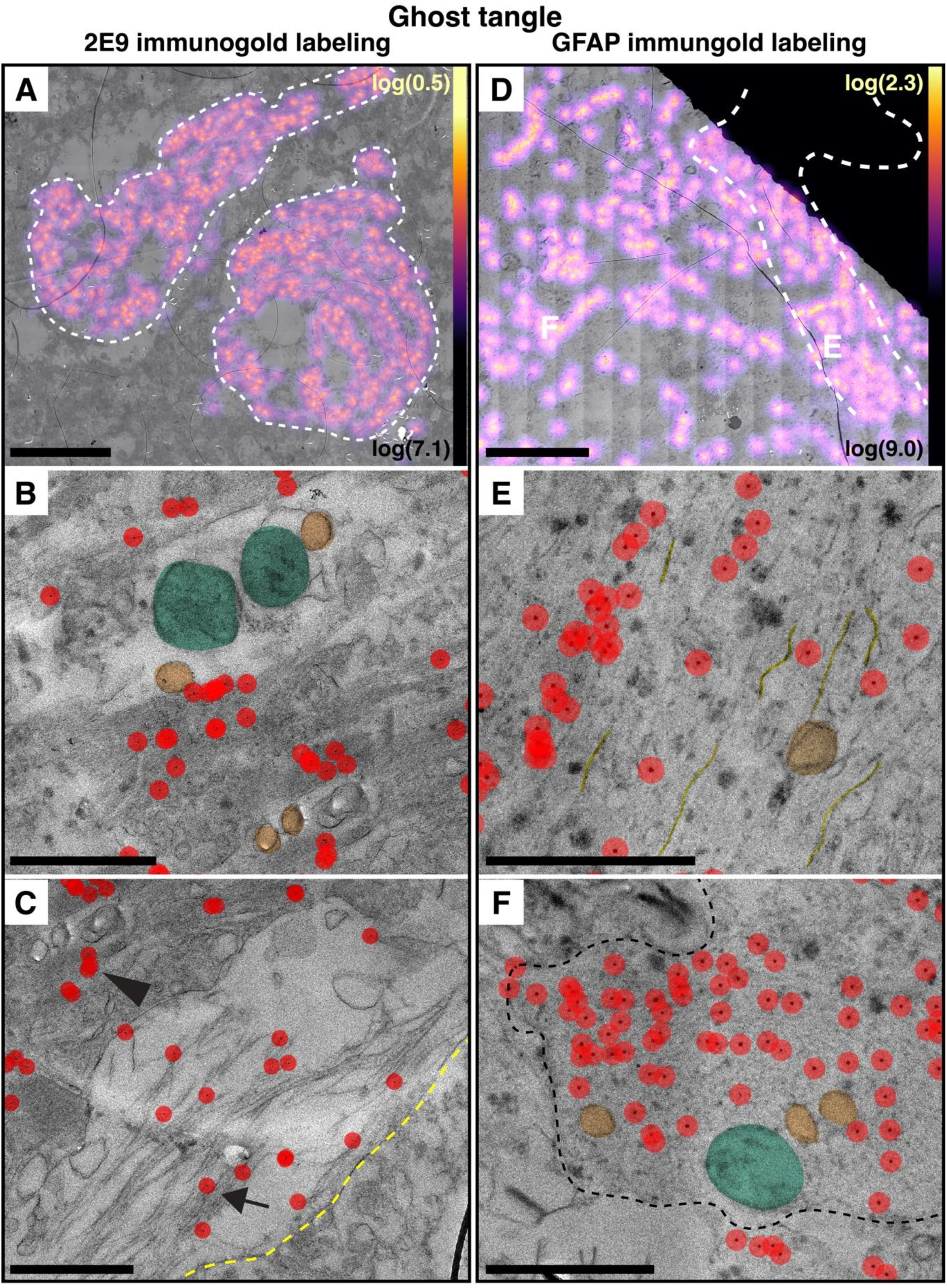
Ghost tangle fibrils are tau-positive by immunogold labelling. **A,** A Low-magnification EM micrograph of two adjacent ghost tangles (white dashed line) immunogold labelled with the fibril core-directed tau antibody 2E9. The heat map indicates the clustering of gold particles over fibril-rich regions. **B,** A high-magnification view showing the enrichment of 2E9 immunogold labelling over thin fibrils and the absence of labelling over vesicles (orange) and mitochondria (green). **C,** 2E9 immunogold labelling of extracellular straight filaments (arrow) and ghost tangle fibrils (arrowhead); areas devoid of fibrils show no labelling (dotted outline). **D,** A low-magnification EM micrograph of a ghost tangle (white dashed line; partially obscured by hexagonal grid bar. This tangle is from an adjacent grid to the one presented in Figure 2A) immunogold labelled with GFAP. The heat map indicates clustering of gold particles over fibrillar regions within the ghost tangle and adjacent astrocytic processes. **E,** A high magnification view showing GFAP labelling within areas of ghost tangle fibrils. Vesicular structures (orange) are unlabeled. Membranes delineating fibrillar compartments, also unlabeled, are indicated (yellow). **F,** GFAP-positive astrocytic process (black dashed line) with labelled filaments. Mitochondria (green) and vesicles (orange) are unlabeled. Scale bars: A, D = 10 µm; B, C, E, F = 1 µm; heat maps in A&D show the summed distance to the five nearest neighboring gold particles for each pixel.

Together, the tomography and immunogold analyses demonstrate that ghost tangles contain a distinct population of tau-immunopositive thin fibrils and are associated with GFAP-positive filaments, distinguishing them from mature tangles, which contain only PHFs and SFs and lack GFAP labeling.

## Discussion

In this study, we performed an ultrastructural analysis of NFT maturation levels in the human AD brain using CLEM. By combining immunofluorescence-based staging with targeted EM, electron tomography, and immunogold labelling, we showed that tau fibrils differ markedly across tangle maturity states. Pre-tangles were devoid of fibrillar material, mature tangles were composed of densely packed bundles of PHFs and SFs amongst morphologically in-tact organelles, and ghost tangles lacked cellular architecture and instead contained compartmentalized assemblies of thinner, non-twisting tau-positive fibrils. Together, these findings establish that tau fibril ultrastructure evolves across NFT maturation and is not static, with distinct structural states emerging during disease progression.

Pre-tangle-bearing neurons, defined by weak p-tau immunoreactivity and absence of β-sheet staining, were ultrastructurally indistinguishable from neighboring p-tau-negative neurons. Consistent with previous ultrastructural studies^12^, organelles appeared intact and no consistent fibrillar structures were observed. Occasional perinuclear filaments were detected but were also present in tau-negative cells, suggesting that these likely represent cytoskeletal elements rather than pathological tau assemblies. These observations support a model in which early tau pathology is primarily molecular, involving the phosphorylation, acetylation and ubiquitination^38^ and mislocalization of tau, and is not yet associated with the formation of stable fibrillar aggregates detectable by EM.

In pre-tangles, we occasionally observed vacuolar structures that may indicate granulovacuolar degeneration, an additional early neuronal response associated with tau pathology (reviewed in^30^). We observed a single structure in a pre-tangle that morphologically resembled a GVB (a vacuole with a dense core), with empty vacuoles observed more frequently. Their identity could not be confirmed due to incompatibility of the tested GVB markers with our glutaraldehyde-fixed tissue, and interpretation is further complicated by the reported variability in GVB abundance between neurons, as well as the unknown effects of post-mortem delay, Braak stage and tissue processing on GVB ultrastructure. Additionally, as Thal GVD stage was not assessed in our cohort, it remains unclear whether the prevalence of GVB-like structures in these donors was unusually low.

In contrast, mature tangles were characterized by large quantities of PHFs and SFs arranged in aligned bundles within the neuronal soma. Despite this substantial fibrillar burden, cellular architecture remained largely preserved, with visible plasma membranes and identifiable organelles including mitochondria, ER and Golgi apparatus. This suggests that extensive fibril accumulation does not immediately result in gross cellular disintegration, consistent with prior reports that neurons can retain substantial ultrastructural integrity despite high intracellular tau load^12,13,18^ and that NFTs may be protective against neuronal death^39^

Mature tangle fibrils were frequently organized into spatially distinct fibril bundles within single neurons, suggesting that tau aggregation may occur at multiple nucleation sites or follow locally constrained growth patterns rather than forming a homogeneous cytoplasmic network. This highly organized intracellular state contrasts sharply with the structural features observed in ghost tangles.

Ghost tangles represented a fundamentally distinct structural state. They lacked a nucleus, neuronal plasma membrane, and intact organelles, and instead consisted of compartmentalized regions of densely packed fibrils delineated by membrane boundaries. These fibrils were consistently thinner, more densely packed, and lacked the characteristic helical twist of PHFs and SFs. Electron tomography confirmed that their diameter was significantly reduced compared to fibrils in mature tangles, while immunogold labelling with the fibril core-directed tau antibody 2E9 demonstrated that they remain tau-positive. Importantly, these fibrils were distinct from astrocytic GFAP fibrils, which were thinner and showed different ultrastructural features.

A defining feature of ghost tangles was the loss of all phospho-tau epitopes tested here and reported previously^6^, despite robust labelling in pre-tangles and mature tangles. These epitopes are in the N- and C-terminal regions of tau that form the fuzzy coat surrounding the amyloid core. In contrast, the tau antibody 2E9, whose epitope maps to the R4 domain of the microtubule binding region falling within the amyloid core of tau fibrils, retained labelling in ghost tangles. As such, the tau fibrils of ghost tangles show a different binding profile with epitope-specific antibodies, consistent with previous reports^7,40^.

Together, the reduction in fibril width, loss of twist, and absence of fuzzy-coat epitopes indicate that ghost tangles do not simply represent residual extracellular PHFs/SFs following neuronal death but instead comprise a structurally distinct population of tau fibrils that emerges at late stages of pathology. These findings suggest that tau aggregates undergo structural remodeling during the transition from intracellular mature tangles to extracellular ghost tangles. One mechanism consistent with our data is proteolytic trimming of the fuzzy coat, preserving the amyloid core and resulting in thinner fibrils. This is supported by the loss of phosphorylation-dependent epitopes and retention of fibril core-directed 2E9 labelling. Alternatively, dissociation of protofilaments may contribute to the reduced fibril width, although this alone does not fully account for the observed changes in morphology and epitope accessibility. These mechanisms are likely not mutually exclusive and may occur in parallel. Consistent with this, previous reports of truncated and less phosphorylated tau species in ghost tangles^41–45^, support the idea that tau fibrils remain susceptible to biochemical processing after neuronal death.

Ghost tangles exhibited a striking compartmentalized organization, in which membrane-bound fibril-rich regions contained GFAP-positive astrocytic processes, consistent with previous reports of astrocyte infiltration of extracellular tau pathology^16–18,33–35^. Notably, the fibrils retained within these compartments, including those associated with astrocytic processes, consistently exhibited the thinner, non-twisting morphology characteristic of ghost tangles, whereas fibrils outside the compartments retained canonical PHF/SF features. These findings raise the possibility that the compartmentalized regions represent sites of astrocytic engagement, in which tau fibrils are engulfed and remodeled through processes such as proteolytic trimming, post-translational modification, or selective degradation of specific fibril species. Although the precise mechanisms remain to be determined, our data suggest that astrocytic engagement may actively contribute to the structural transformation of tau during ghost tangle formation.

Collectively, our observations support a stage-dependent model of tau aggregation, in which fibril architecture and composition evolve during disease progression, rather than remaining static. In this model, early tau pathology is dominated by non-fibrillar tau species, followed by the formation of highly ordered PHF and SF bundles within intact neurons. At later stages, these assemblies transition into extracellular ghost tangles, where tau fibrils undergo structural remodeling, resulting in thinner, non-twisting filaments within a compartmentalized environment. The compartments include infiltration by surrounding astrocytic processes, which may contribute to fibril remodeling (**Fig. 5**).

**Figure 5:**
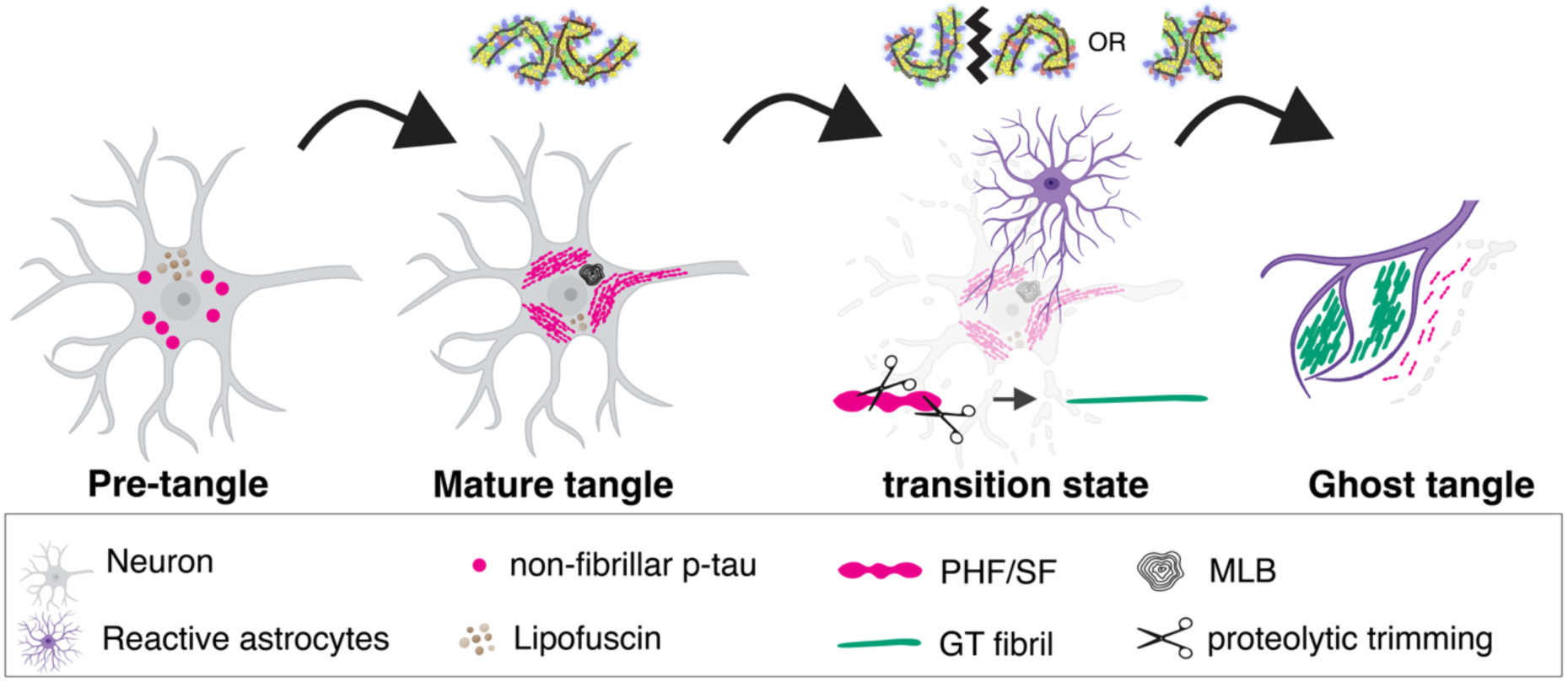
Model of tau fibril formation and remodeling across NFT maturation stages. A schematic representation of tau aggregation across pre-tangles, mature tangles, and ghost tangles. Pre-tangles contain little to no detectable fibrillar material and are likely dominated by non-fibrillar p-tau species. In mature tangles, multiple nucleation sites give rise to the progressive accumulation of PHFs and SFs, which form aligned, densely packed bundles within a largely preserved neuronal soma. Following neuronal death, tau aggregates transition into ghost tangles. During this stage, astrocytic processes infiltrate and compartmentalize fibrillar material. The tau fibrils^46,47^ undergo structural remodeling, potentially through protofilament separation and/or proteolytic trimming of N- and C-terminal regions. Fibrils located in what appears to be the extracellular space retain features of canonical PHFs/SFs, whereas fibrils within astrocyte-associated compartments appear thinner and lack the characteristic helical twist. This figure was partially made with BioRender Lewis, A. (2026) https://BioRender.com/wmohscv.

These EM findings have biological and therapeutic implications. First, they indicate that tau aggregates are not terminal end-products of fibrillization but continue to evolve structurally at late stages of disease. Second, the loss of phosphorylation-dependent epitopes in ghost tangles has implications for biomarker development and imaging strategies, many of which rely on p-tau detection and may therefore underestimate late-stage pathology. Third, the apparent susceptibility of tau fibrils to astroglial remodeling raises the possibility that endogenous pathways of fibril processing could be therapeutically targeted to enhance clearance or modify pathogenic species.

Resolving the molecular architecture of ghost tangle fibrils will be critical to elucidate these mechanisms. However, this remains technically challenging, as no antibodies currently available selectively target ghost tangle fibrils, and their behavior under conventional fibril purification protocols, such as sarkosyl treatment, is unknown. Recent advances in cryo-CLEM and in situ cryo-electron tomography provide a promising route to address this question. Volume EM studies would help to further characterize the astrocytic infiltration of mature tangles and their transition to ghost tangles. While recent studies have resolved tau fibril structures directly within human brain tissue^21^, our observations indicate that most fibrils in ghost tangles are structurally distinct from canonical PHFs and SFs, suggesting that current structural datasets does not capture the diversity of tau assemblies present at late stages of disease.

Our study has several limitations. The low throughput nature of CLEM restricted our analysis to a limited number of donors and brain regions with advanced pathology and therefore does not address regional or disease-stage variability. While we demonstrate that ghost tangle fibrils are tau-positive and distinct from GFAP, the resolution of room-temperature CLEM does not allow determination of atomic structure or isoform composition^22^. Furthermore, as this study is based on post-mortem human tissue, it provides static snapshots and cannot resolve the temporal sequence of tangle maturation or astrocytic involvement.

In summary, our CLEM analysis reveals that tau pathology undergoes substantial structural and compositional changes during neurofibrillary tangle maturation in the human AD brain. While mature tangles consist of canonical PHF and SF bundles within a preserved neuronal environment, ghost tangles contain thinner, structurally distinct tau fibrils organized within a compartmentalized extracellular context. These findings demonstrate that tau aggregates are dynamic structures that continue to remodel at late stages of disease and provide a framework for understanding the evolution of tau pathology with implications for biomarker interpretation and therapeutic intervention.

## Materials and Methods

### Human post-mortem AD brain samples

In this study, postmortem brain tissue was collected from four AD cases (Donors A– D) enrolled in the Netherlands Brain Bank (NBB) donation program (www.brainbank.nl), each with a post-mortem delay under 6 hours (**Suppl. Table 1**). At autopsy, the NBB’s standardized dissection protocol was used to isolate blocks from the hippocampus and entorhinal cortex for all donors. At autopsy, subregions of the hippocampus (CA1 and CA2), entorhinal cortex and trans-entorhinal cortex were dissected from Donor A and B into small cubes of 2 mm^3^ and fixed for 24 hours in 4% paraformaldehyde (PFA) and 0.1% glutaraldehyde in 0.15 M cacodylate buffer (pH 7.4), supplemented with 2 mM calcium chloride. Tissues of Donor C and D were fixed in 4% PFA at autopsy and post-fixed overnight at 4 °C in 4% PFA with 0.1% glutaraldehyde in cacodylate buffer before processing for CLEM. All donors had provided written informed consent for brain autopsy and the use of their tissue and clinical data for research and all procedures were performed in accordance with the declaration of Helsinki. Detailed neuropathological diagnoses and relevant clinical information were obtained in accordance with local ethical and legal guidelines, and all procedures were approved by the Amsterdam UMC institutional review board (reference 2019/48; www.brainbank.nl). Demographic and clinical data—including sex, age at symptom onset, age at death, disease duration, presence of dementia, and other core or supportive clinical features—were extracted from medical records (**Suppl. Table 1**).

For pathological diagnosis, 7-μm-thick formalin-fixed paraffin-embedded sections were immuno-stained using antibodies against αSyn (clone KM51, 1:500, Monosan Xtr), amyloid-β (clone 4G8, 1:8000, Biolegend) and phosphorylated tau (p-tau, clone AT8, 1:500, ThermoFisher Scientific), as previously described^48^. Braak and McKeith αSyn stages were determined using the BrainNet Europe (BNE) criteria^49^. Thal amyloid-β phases were scored on the medial temporal lobe^50^, Braak neurofibrillary stages^2^ and Consortium to Establish a Registry for Alzheimer’s Disease (CERAD) neuritic plaque scores^51^, levels of Alzheimer’s disease (AD) pathology were determined according to National Institute on Aging-Alzheimer’s Association (NIA-AA) consensus criteria^52^. Additionally, Thal cerebral amyloid angiopathy (CAA) stages^44^, presence of ageing-related tau astrogliopathy (ARTAG)^9^, micro-vascular lesions and hippocampal sclerosis were assessed (**Suppl. Table 1**).

### CLEM

Correlative Light and Electron Microscopy (CLEM) studies were performed as we previously described^22^ with the following changes.

### Immunofluorescence staining

Briefly, 20 - 60 μm thick brain sections were collected with a vibratome (Leica VT1200). Free-floating sections were immunolabelled with one anti tau antibody (pT217; Thermo Fisher Scientific #44-744, 1:500, AT8; Thermo Fisher Scientific #MN1020, 1:200, or AT180; Thermo Fisher Scientific #MN1040 1:500), a cell nuclei marker (DAPI; Biolegend #422801, 1:800) and a β-sheet marker (Amytracker 480; Ebba Biotech #A480-A-50, 1:500-1:1000). Sections were incubated with the primary anti-tau marker at 4°C, ON followed by incubation with Alexa-conjugated secondary antibodies, DAPI and Amytracker for 30 minutes at room temperature. The sections were mounted on glass slides in 50% TBS-glycerol for fluorescent imaging.

### Heavy metal staining and resin embedding

Sections were processed using a Leica Tissue Autoprocessor (Leica model EM TP). Sections were postfixed in 2% osmium tetroxide, reduced with 0.15% (wt/vol) potassium ferrocyanide, fixed again in 2% osmium tetroxide and stained in 2% uranyl acetate for 30 minutes. To enhance contrast, sections were additionally stained with lead aspartate, pH 5.5 at 60°C. Sections were dehydrated in a graded ethanol series and embedded in EMBED 812 resin. Serial sections of 80–200 nm were cut using an ultramicrotome (Leica, EM UC7) and alternatingly collected on electron microscopy grids and glass slides, respectively.

### Immunohistochemistry

150 - 200 nm semi-thin resin sections on glass slides were processed for immunohistochemistry using antibodies against tau and GFAP. Sections were first etched in saturated potassium ethoxide for 3 mins before the addition of the primary antibody. The following conditions were used for the individual antibodies: AT8 and AT180 were used at 1:1000, 4h at RT. The anti tau antibody 2E9 (NovusBiological # NBP2-25162) was used at 1:2000, 1h, 37°C. For 2E9, antigen retrieval was needed, and glass slides were first incubated with 100% formic acid, 10 min at RT followed by steaming in Tris-EDTA, pH 9 for 30 min at 100°C. Anti-GFAP antibody AB5531 (Merck #AB5541) was used at 1:1000, 1 h at 37°C. For AB5531, antigen retrieval was required as well and glass slides were incubated with 100% formic acid, 10 min at RT followed by steaming in either Tris-EDTA, pH 9 or Citrate buffer, pH 6 for 30 min at 100°C. ImmPRESS Reagent anti-mouse or anti-rabbit secondary antibodies were used (Vector Laboratories) for 30 min at room temperature. Bound antibody complexes were detected using the permanent horseradish peroxidase (HRP) Green Kit (Zytomed Systems) with incubation for 3 min at RT or the ImmPACT^®^ NovaRED^®^, Substrate Kit Peroxidase (Vector Laboratories) with incubation for 5 min at RT. Sections were counterstained with either haematoxylin or toluidine blue and mounted on glass coverslips for imaging.

### Fluorescent image correlation

Images were correlated based on tissue features such as blood vessels and cell nuclei or in some cases, introduced laser marks. For making the overlays, we used the FIJI plugin FIJI BigWarp^53,54^. Pathologies themselves were helpful for orientation, however, were typically not used for image correlation. Fluorescent images were correlated to the IHC images, which served as an intermediate reference for direct correlation with EM images.

For NFT maturity level characterization, fluorescent, IHC and EM images were used. While with immunofluorescence it was not always clear if a nucleus was present, this became apparent in IHC and EM images. To distinguish between dystrophic neurites and mature tangles, we checked for the presence of a cell nucleus or soma-related cell organelles as the Golgi apparatus or endoplasmic reticulum. In some cases, we could also follow the same pathology through several CLEM cycles and confirm the presence of a nucleus in a different cycle. A CLEM cycle consisted of two or three 200 nm sections per glass slide and two or three 80 nm sections per EM grid, corresponding to a total depth of ca. 1.5 μm. Sections were collected on 2 separate glass slides, which allowed two independent IHC experiments, and on two copper slot grids and one hexagonal grid (FCF2010-CU-TA-50ST, FF-100H-Cu-UL; Electron Microscopy Sciences). Typically, 10 cycles were collected per block, corresponding to ca. 15 μm in depth.

### Immunogold labelling

Immunogold labelling was performed using tau (2E9) or GFAP (AB5531) antibodies on the EM grids based on the standard protocol provided by Aurion. Briefly, the resin was etched for 3-10 min in 1% periodic acid (Sigma). Mild antigen retrieval was performed by incubating EM grids with Tris-EDTA, pH 9 for 1 hour at 37°C. Other conditions, such as using formic acid, or steaming at 100°C were found to damage the samples on the grids, so that these were not pursued. Since sections placed on slot grids often broke during immunogold labeling, hexagonal grids were used. As a test, grids that had already been imaged by the EM, were subjected to immunogold labeling, but this labeling failed on the pre-imaged areas, possibly due to the presence of electron beam damage to protein structures. For secondary labeling, grids were incubated with 10 – 15 nm protein A conjugated gold beads (Aurion) or 10 nm donkey anti-mouse antibody conjugated gold beads (Aurion) for 90 min at room temperature. To reduce contamination and confounding results, re-contrasting with uranyl acetate was omitted. Several individual EM montages were recorded for ghost tangles and mature tangles, and different GFAP immunogold positive areas were imaged. Gold beads were detected using a custom FIJI macro and python notebook (https://github.com/lukasvandenheuvel/dog-immunogold-detection), and heat maps were generated by calculating pixel-wise the average distances to closest 5 neighboring gold beads.

### Light Microscopy

Features in the tissue that were identifiable in both the light and electron microscopy images were used to guide the collection of EM micrographs. Fluorescent images were collected on a Leica Thunder light microscope, equipped with a Leica K8 camera and a colour Leica FLEXACAM-C1. The light sources and filter systems had the following properties (excitation/emission): 395/440; 480/510; 555/590; 640/700. Fluorescent images were recorded either with a HC PL APO 10x, 0.45 NA air objective (for creating overview maps) and with a HC PL APO 40x, 0.95 NA air objective or a HC PL APO 63x, 1.40 NA oil objective for higher resolution z-stacks of pathologies of interest. Z-stacks at higher resolution were typically collected at 1 μm step size. All 4 available channels were recorded. The microscope’s yellow channel was not used to avoid confounding real staining with background (mostly stemming from lipofuscin, which emits mostly in the 480 and 555 channels). Fluorescent images were processed and deconvoluted using the Leica Thunder algorithm^55,56^. IHC bright field images were recorded using the HC PL APO 63x, 1.40 NA oil objective. Consecutive image tiles were stitched together using the Leica LAS X software.

### Electron Microscopy

Micrographs were typically collected at a pixel size of 1.7-1.9 nm/pixel at room temperature. Images were recorded with a Philips CM100 Biotwin TEM operated at 80 kV with a LaB6 filament electron source and a bottom-mounted TVIPS F416 camera, or a FEI Tecnai Spirit BioTwin (120kV) with a LaB6 filament and a bottom-mounted TVIPS F416 camera, operated at 80 kV for section imaging or at 120 kV for tomography data collection, or a ThermoFisher Scientific (TFS) TALOS (120 kV) equipped with a with TFS CETA camera operated at 120 kV.

### Image Display

In some cases, images were filter adjusted using BaSiC and Rolling Ball plugins in FIJI^54,55^. Signal from the 640/700 channel was represented in magenta instead of red to improve visibility for colour-blind readers, and both light and electron microscopy images were adjusted for brightness and contrast where necessary using FIJI.

### Tomography

Tilt series were collected either at the Tecnai equipped with bottom mount TVIPS F416 camera or at a Talos equipped with a CETA camera operated at 120 kV at RT at a pixel size of 0.5 nm. Tomography data were collected with SerialEM^56,57^ on the Tecnai and TFS Tomography 5 on the Talos and collected images at −60° to +60° tilt angles with a 2° step size. Tomograms were binned by a factor of 2 and filtered using a non-local means filter in Amira version 2021.2 (TFS). Segmentation of the fibrils was carried out using the EMAN2 semi-automated convolutional neural network (CNN) protocol^58^ and refined using the UCSF Chimera package^59^ and Amira. The fibril thickness distribution was extracted using the Amira Thickness Map module. The extracted data were thresholded to match the EMAN2 segmentation and to exclude overlapping fibrils when crossing each other. Fibril widths were measured with a line tracer.

### Statistical Analysis

Each dataset was screened for outliers by calculating the Z-score for every individual measurement within its respective group. Measurements with Z-scores below −3 or above 3 were considered outliers and excluded from further analysis. The distribution of the remaining data was then assessed using the Shapiro–Wilk test to verify the assumption of normality. As all datasets were normally distributed after outlier removal (*p* > 0.05), group comparisons were performed using a two-tailed Welch’s *t*- test assuming non-equivalent standard deviation. The Welch’s *t*-test was chosen because it does not require equal variances between groups. Differences were considered statistically significant at *p* < 0.05.

Statistical analyses were performed using GraphPad Prism 11 or MicExcel.

## Data availability

All raw EM images will be uploadedto the BioImage Archive (https://www.ebi.ac.uk/bioimage-archive/) upon acceptance of this manuscript for publication.

## Author Contributions

DAS, HS & AJL designed the study. DAS & LT performed the CLEM, immunogold labelling and electron tomography, with assistance from EV, PLS, LvdH and NS. NS performed statistical analysis. AR and WVdB performed the brain dissection and neuropathological evaluation. DAS prepared the figures. DAS and AJL wrote the manuscript. All authors assisted with data interpretation and approved the final version of the manuscript.

## Acknowledgements

We are deeply grateful to the brain donors and their families whose generous participation in the brain donation program made this work possible. We thank all members of the Netherlands Brain bank autopsy team for the provision of high-quality post-mortem brain tissue for electron microscopy. We also appreciate the technical support and upkeep provided by the Electron Microscopy Facility staff at the University of Lausanne. We thank Melissa E. Murray and Christina M. Moloney for their feedback and advice on the characterization of tau tangle maturity levels.

