## Supplementary information for "Ultrastructural remodelling of tau fibrils during ghost tangle formation in Alzheimer’s disease brain"

| Donor-ID | Sex (m/f) | Age (y) | Thal (x/5) | Braak (x/6) | ABC | CERAD (x/3) | TDP | CAA | ARTAG |
|---|---|---|---|---|---|---|---|---|---|
| A | f | 75 | 5 | 5 | 3 3 3 | 3 | No | Type2, 1/3 | no |
| B | m | 70 | 5 | NA | 3 3 3 | NA | NA | Type1, 2/3 | yes |
| C | f | 86 | 5 | 4 | 3 2 2 | NA | No | No | yes |
| D | m | 77 | 5 | 5 | 3 3 3 | 3 | No | Type 1, 3/3 | yes |

**Supplementary Table 1: Demographics and pathological characteristics of included AD donors.** m/f=male/female; y=years; PMD = post-mortem delay; Clin.=clinical diagnosis; neuropat.=neuropathological diagnosis; ABC=ABC criteria according to Montine et al.; CERAD = Consortium to Establish a Registry for Alzheimer's Disease; CAA=Cerebral Amyloid Angiopathy; ARTAG=Aging-related tau astrogliopathy.

| Tau maturity | Clone/name | Target | Fluorescence | IHC | Immunogold | Manufacturer | Product # |
|---|---|---|---|---|---|---|---|
| GVB, PT | sc-55553 | casein kinase 1delta (C-8) | not working | not tested | not tested | Santa Cruz Biotechnology | sc-55553 |
| | sc-373912 | casein kinase 1epsilon (A-2) | not working | not tested | not tested | Santa Cruz Biotechnology | sc-373912 |
| PT, MT | 44-744 | pT217 | 1:500 | not tested | not tested | Thermo Fisher Scientific | 44-744 |
| | AT8 | pS202; pT205 | 1:200 | 1:1'000 | 1:20 | Invitrogen | MN1020 |
| | ERP2400 | pS198 | 1:500 | not tested | not tested | Abcam | ab79540 |
| | 2H23L4 | pS199 | 1:500 | not tested | not tested | Thermo Fisher Scientific | 2H23L4 |
| | E178 | pS396 | 1:500 | not tested | not tested | Abcam | ab32057 |
| | 44-752G | pS396 | 1:500 | not tested | not tested | Thermo Fisher Scientific | 44-752G |
| | E178 | Tau (pS396) | 1:500 | not tested | not tested | Abcam | ab32057 |
| | AT180 | Tau (pThr231) | 1:500 | 1:1'000 | not tested | Thermo Fisher Scientific | MN1040 |
| | AT100 | Tau (pThr212, pSer214) | not working | not tested | not tested | Thermo Fisher Scientific | MN1060 |
| MT, GT | ghost-tangle-38 | Tau conformer | not working | not tested | not tested | Abcam | AB246808-1001 |
| | 2E9 | aa 347-366 | 1:500 | 1:1'000 | 1:50 | Novus biologicals | NBP2-25162SS |
| | TauC3 | aa 412-421 (C-truncated) | not working | not tested | not tested | Santa Cruz Biotechnology | sc-32240 |
| | Amytacker 480 | β-sheets | 1:1'000 | - | - | Ebba Biotech | #A480-A-50 |

**Supplementary Table 2: List of tested antibodies and markers used in this study.** Working concentrations are indicated. Granulovacuolar bodies (GVB); Immunohistochemistry (IHC); Pre-tangle (PT); Mature tangle (MT); Ghost tangle (GT).

| Patient | A | B | C | D | Total |
|---|---|---|---|---|---|
| # Pretangles | 7 | 0 | 1 | 0 | 8 |
| # Mature tangles | 28 | 15 | 0 | 0 | 43 |
| # Ghost Tangles | 0 | 40 | 0 | 1 | 41 |

**Supplementary Table 3: Number of pathologies found in this study per patient.**

The counts reported here reflect only the pathologies sampled for correlative analysis and do not represent the overall pathological burden in the donor tissue or patient. Pathologies were identified either by 1) correlation of fluorescence images to EM images, 2) correlation of IHC images to EM images or 3) by ultrastructural assessment or by a combination of these 3 assessment methods. All pre-tangles and ghost tangles presented in this study were characterized either by fluorescence or IHC or both.

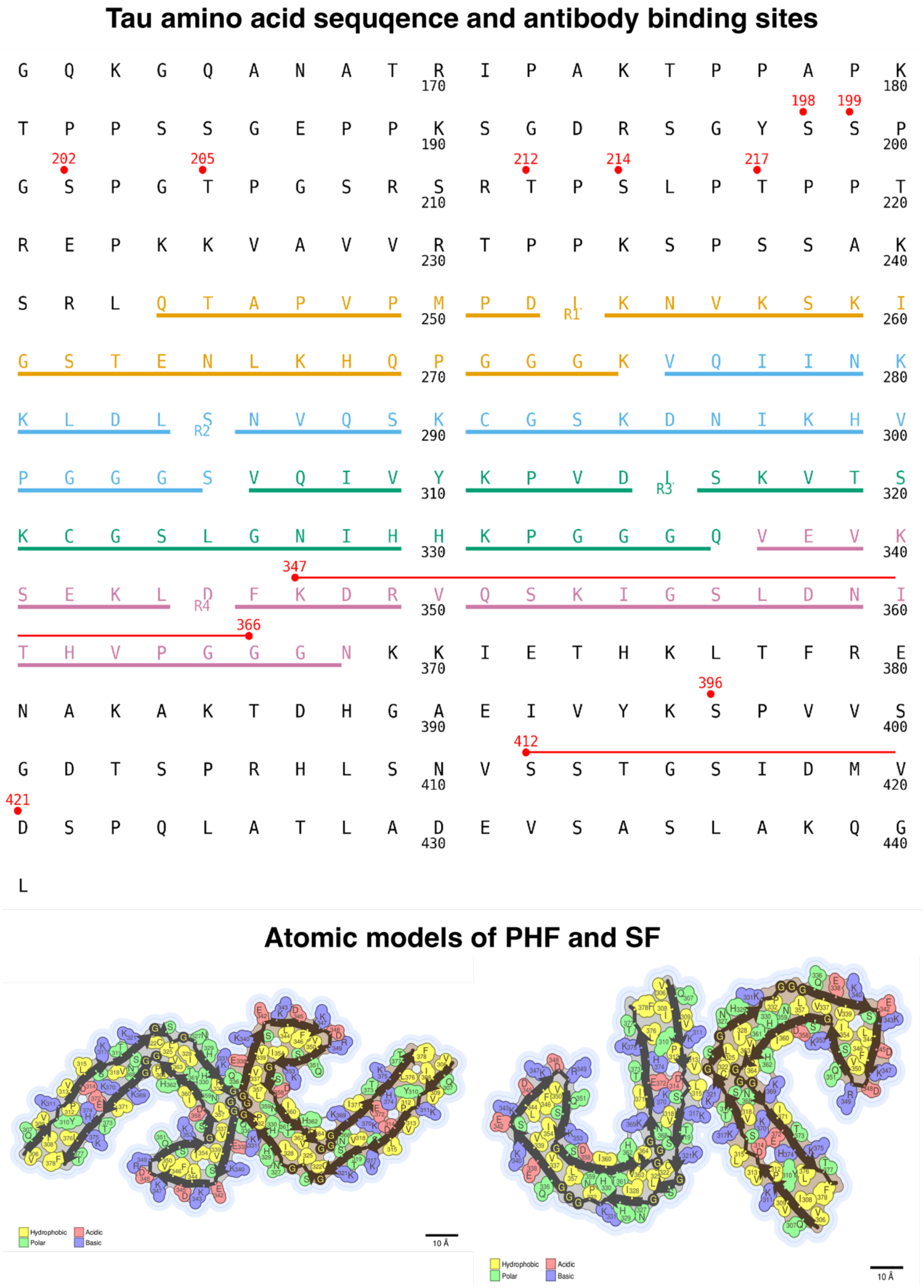


**Supplementary Figure 1: Tau sequence, repeat domains and fibril structures**

The amino acid sequence of human tau (2N4R) with repeat domains (R1-R4) color-coded. The binding epitopes of antibodies tested in this study are indicated (red dots/lines). Note that all binding sites are outside of the R domains, except for 2E9 which binds between aa 347 and 366. Polarity maps of PHFs[1] (left), and SFs[2] (right) encompass aa 306-378 and are derived from cryo-EM structures of tau fibrils extracted from the human brain of AD patients. Polarity maps were made with UCLA's amyloid illustrator[2].

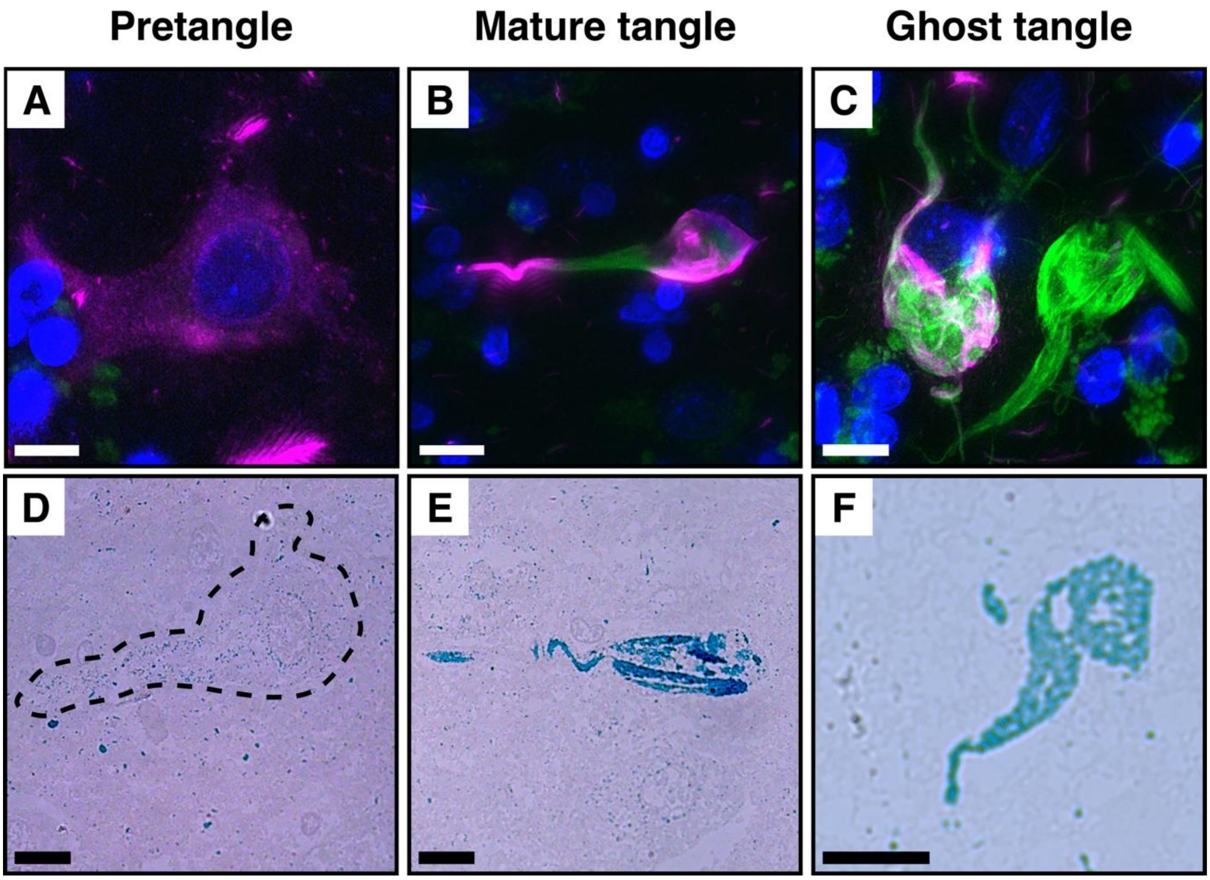


**Supplementary Figure 2: Characterization of NFT maturity states by fluorescence and IHC. A-C,** Fluorescence images showing phosphorylated tau (pT217, magenta), β-sheet signal (Amytracker, green) and nuclei (DAPI, blue). **A,** Pre-tangles display weak diffuse p-tau staining with a nucleus and minimal Amytracker signal. **B,** Mature tangles are positive for p-tau and Amytracker (partially overlapping) and associated with a nucleus. **C,** Ghost tangles show strong Amytracker signal without p-tau and lack an associated nucleus. **D-F,** Corresponding IHC on resin sections. **D,** Pre-tangles show weak dotted p-tau staining (AT8) outlining the neuron. **E**, Mature tangles show strong tau staining (2E9) extending into neuronal processes with a visible nucleus. **F,** Ghost tangles also show strong tau staining (2E9) but lack a nucleus. Scale bars: A, D–F = 10 μm; B = 100 μm; C = 20 μm.

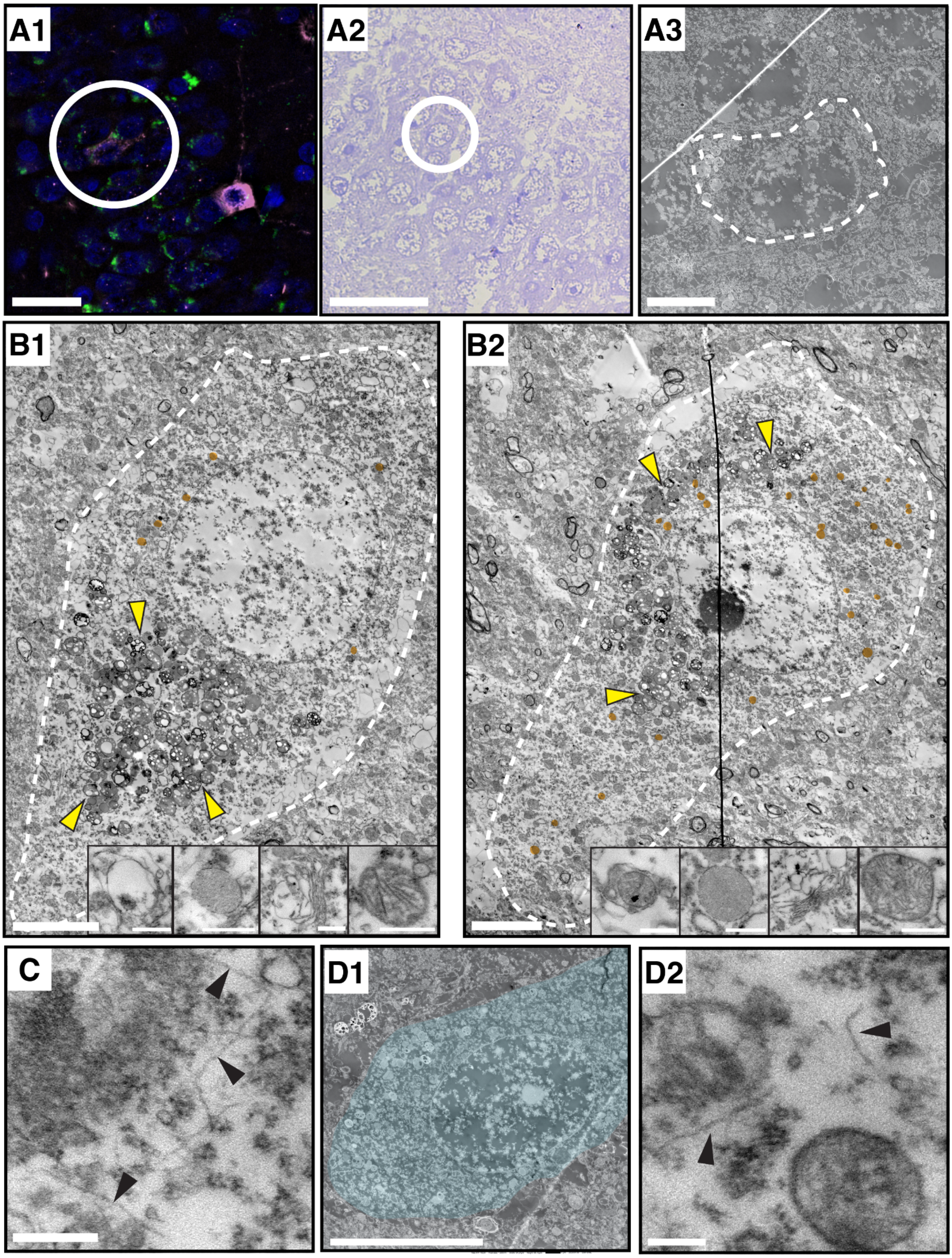


**Supplementary Figure 3: Pre-tangle are ultrastructurally similar to tau-negative neurons. A1–A3,** Correlative fluorescence, toluidine blue and EM of a pre-tangle (outlined). No consistent ultrastructural differences are observed compared with neighbouring cells. **B1,B2** Pre-tangles showing lipofuscin (yellow arrowheads) and lysosomes (orange). Insets show the ultrastructure of a suspected GVB-like structure, defined as a vacuole with a dense core, together with lysosomes, Golgi apparatus and a mitochondria (in B1) and an autophagosome, lysosomes, Golgi apparatus and mitochondria (in B2) respectively. Abundant lysosomes were observed in B2. **C,** Perinuclear fibrils in a different pre-tangle. **D1,D2,** Tau-negative neuron showing intact organelles, lipofuscin and cytoskeletal filaments. Fibril-like profiles similar to those in pre-tangles are occasionally present (arrowheads). Scale bars: A1, A2 = 30 µm; A3 = 5 µm; B1,B2 low mag = 5 µm, insets = 500 nm; C = 300 nm; D1 = 10 µm; D2 = 200 nm.

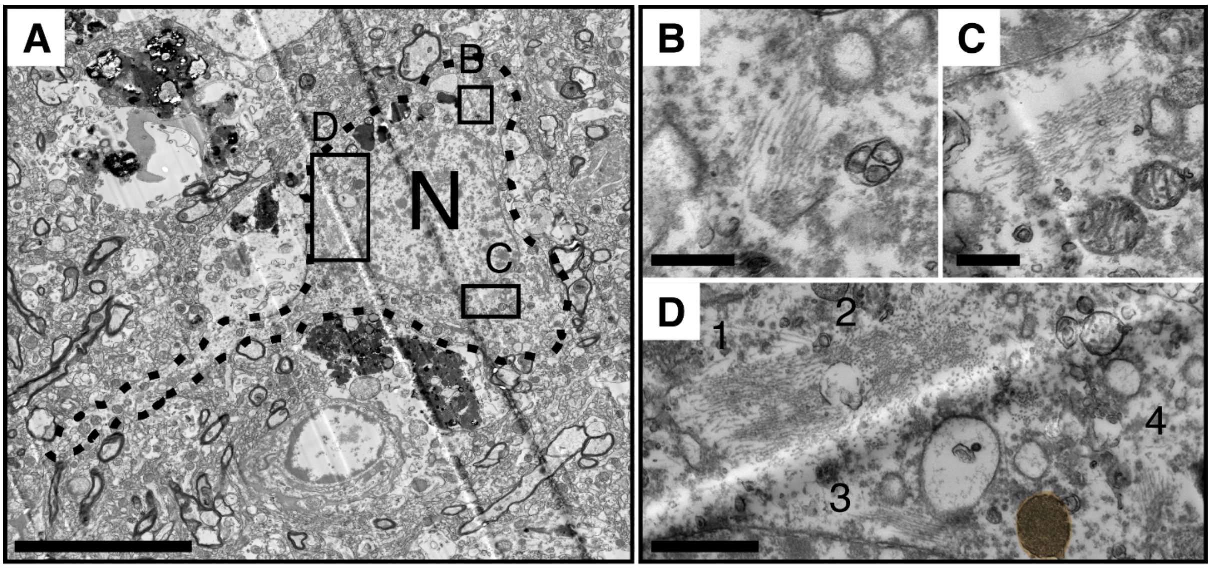


**Supplementary Figure 4: A mature tangle containing spatially separated tau fibril clusters. A,** An overview of a mature tangle showing the soma and neuronal process (outlined). **B-D**, Multiple aligned-fibril bundles showing separation within the cytoplasm. Bundles can differ in orientation relative to the section plane, indicating distinct local organization. **D,** Four distinct clusters are visible, indicated by numbers. Clusters 1 and 2 are in close proximity to each other, and while fibrils in cluster 1 lie parallel to the section plane, fibrils in cluster 2 are oriented orthogonally to the section plane. A single granular lysosome was observed in this neuron (coloured orange.) Scale bars: A = 10 μm; B, C = 500 nm; D = 1 μm.

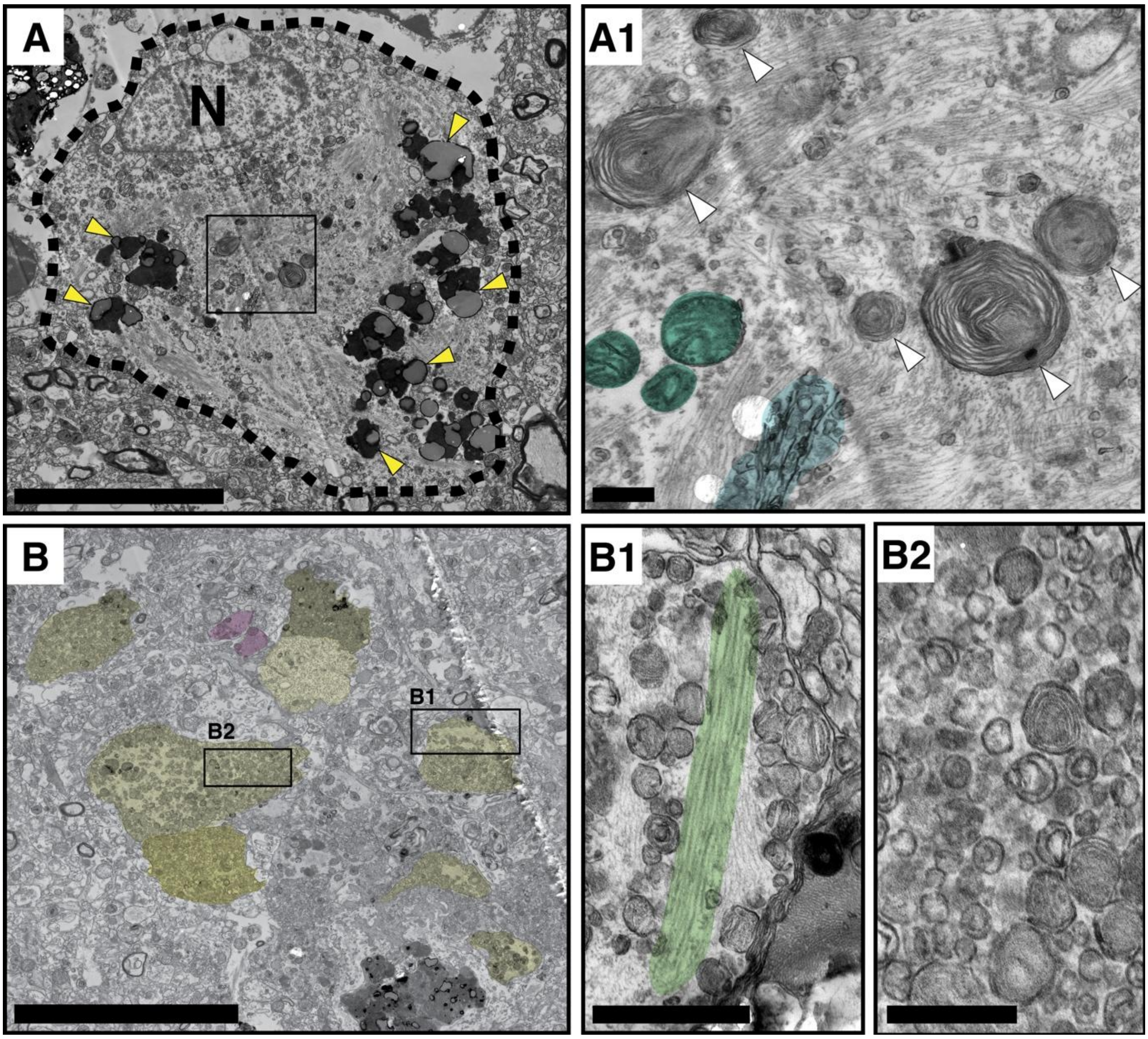


**Supplementary Figure 5: Example of a mature tangle and dystrophic neurites around an Aβ plaque. A, A1,** A mature tangle filled with tau fibrils and lipofuscin (yellow arrowheads) with the nucleus (N) visible in the same plane, with a higher magnification shown in a1. Multilamellar bodies (white arrowheads), mitochondria (green), Golgi (blue) and vesicular membranes are visible between the fibrils. **B,** Dystrophic neurites (yellow) and neuropil threads (magenta) surrounding the tangle, with higher magnifications shown in b1 and b2. **B1,** Some dystrophic neurites contain tau fibrils (green). **B2,** Other neurites are dominated by multilamellar and multivesicular structures. Scale bars: A, B = 10 µm; A1, B1 = 1 µm; B2 = 500 nm.

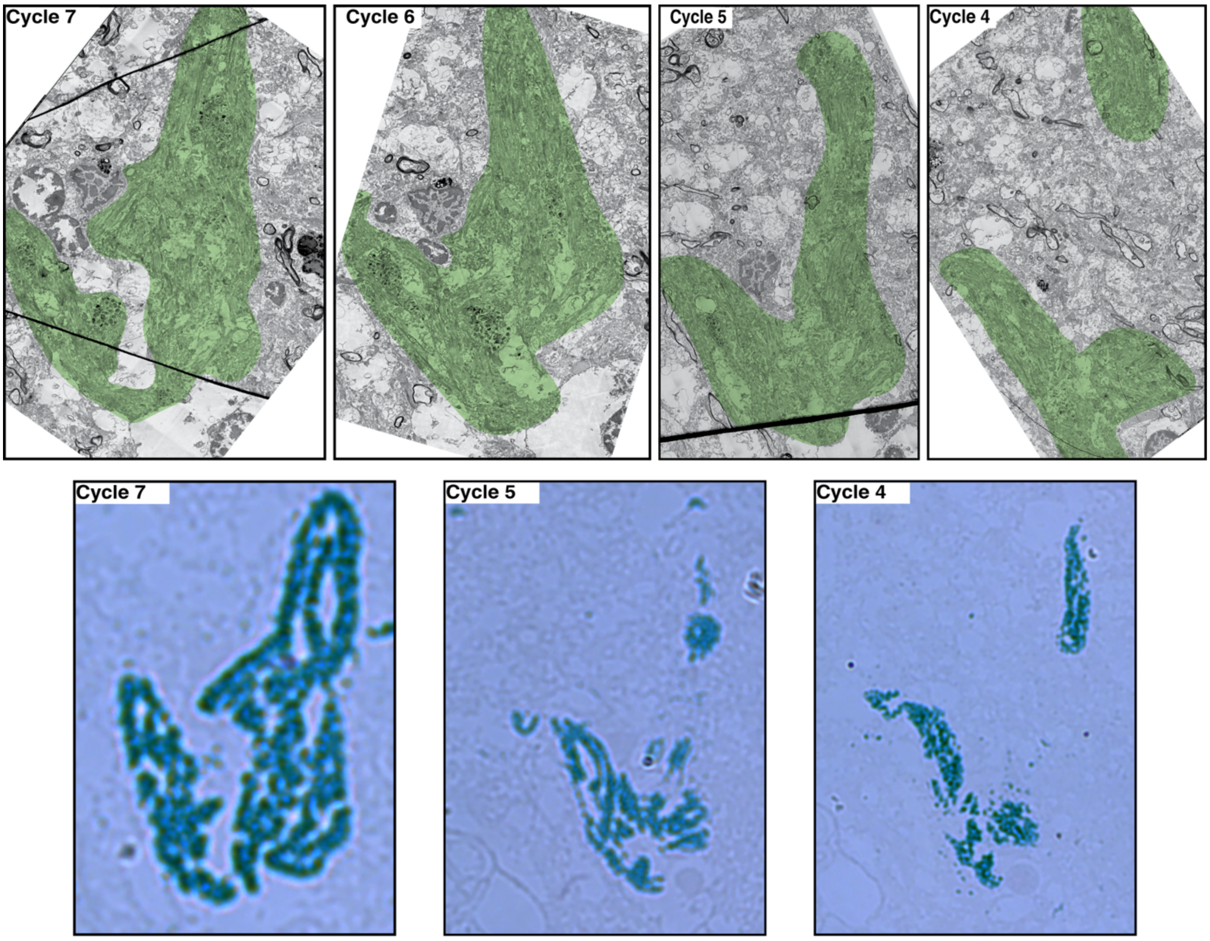


**Supplementary Figure 6: Ultrastructure of a ghost tangle across several CLEM cycles.** Serial EM images of the same ghost tangle across successive CLEM sections. Thin fibril-rich regions (green overlay) correspond to 2E9 tau immunoreactivity on adjacent IHC sections. The compartmental organization is preserved throughout the analyzed depth.

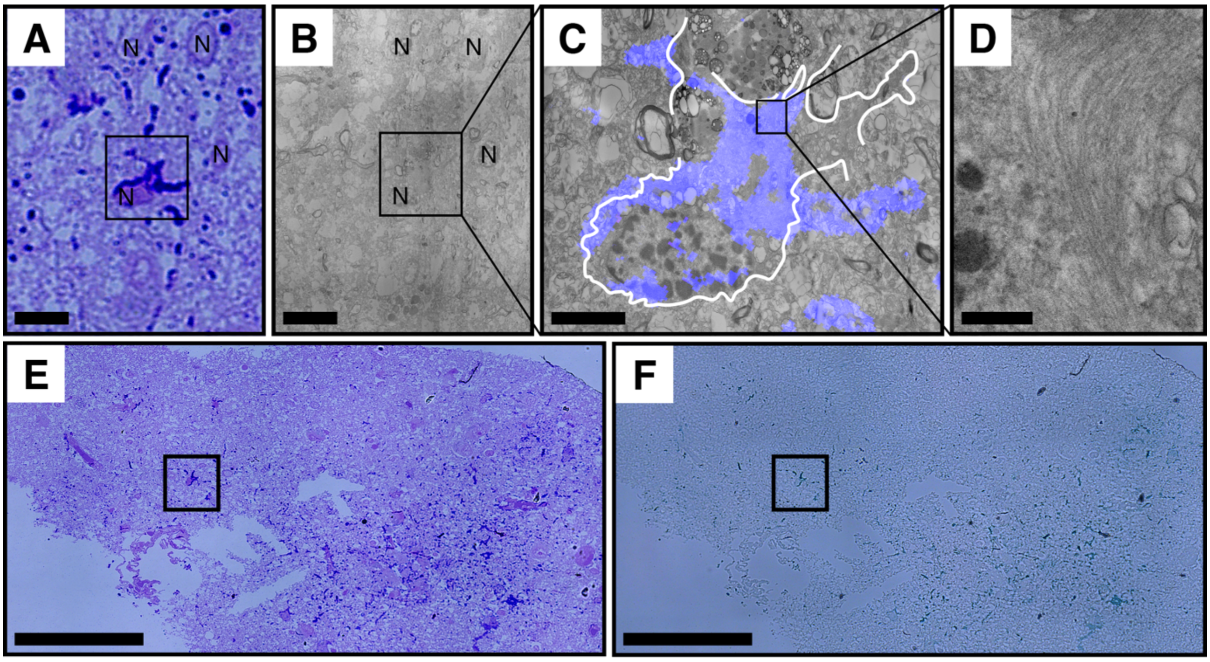


**Supplementary Figure 7: IHC and ultrastructure of astrocytic GFAP filaments. A,** IHC of an astrocyte labelled with anti-GFAP showing the cell body and processes. **B,C,** The correlated EM regions. **D,** A high magnification view of the densely packed GFAP filaments showing the characteristic thin, non-twisting, curved morphology distinct from PHFs/SFs. **E,F,** Overview of the tissue section before and after staining enhancement by toluidine blue. Scale bars: A, B = 10 μm; C = 3 μm; D = 300 nm; E, F = 100 μm.

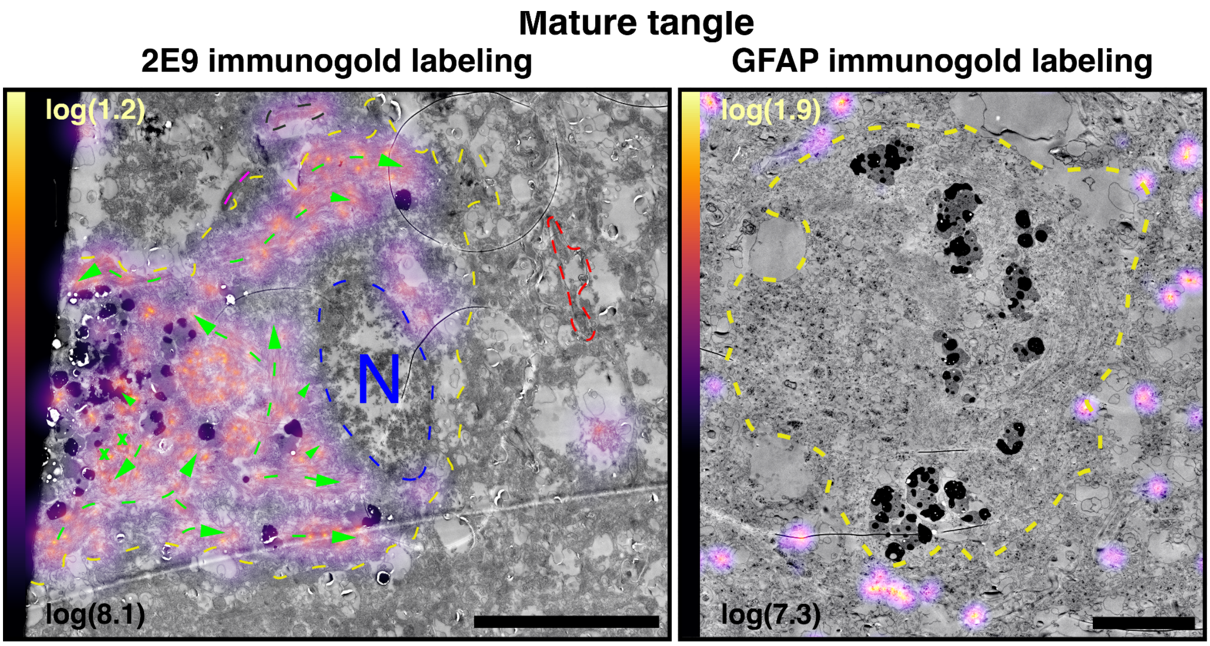


**Supplementary Figure 8: Mature tangle fibrils stain positive for 2E9 tau marker but not for the anti-GFAP marker using immunogold labeling. A,** 2E9 immunogold labelling shows enrichment over tau fibrils within a mature tangle (yellow dashed line) and a neuropil thread (black dotted line), with no labelling of adjacent astrocytic processes (pink dotted line) or axons (red dashed line). A nucleus is visible (N) within the tangle-bearing neuron, and the presence and direction of fibrils is indicated (green lines with arrows). **B,** GFAP immunogold labelling is restricted to astrocytic processes outside the mature tangle (yellow dashed line) and does not label tau fibrils. Scale bars: A = 10 µm; B = 5 µm; Heat maps show the summed distance to the ten nearest neighboring gold particles for each immunogold particle.
